# A practical guide to CRISPR/Cas9 genome editing in Lepidoptera

**DOI:** 10.1101/130344

**Authors:** Linlin Zhang, Robert D. Reed

## Abstract

CRISPR/Cas9 genome editing has revolutionized functional genetic work in many organisms and is having an especially strong impact in emerging model systems. Here we summarize recent advances in applying CRISPR/Cas9 methods in Lepidoptera, with a focus on providing practical advice on the entire process of genome editing from experimental design through to genotyping. We also describe successful targeted GFP knockins that we have achieved in butterflies. Finally, we provide a complete, detailed protocol for producing targeted long deletions in butterflies.

## Introduction

The order Lepidoptera represents a tenth of the world’s described species and includes many taxa of major scientific and agricultural importance. Despite strong interest in this group, however, there has been a frustrating lack of progress in developing routine approaches for manipulative genetic work. While the last decade has seen examples of transgenesis and targeted knockouts using methods like transposon insertion^1^, zinc-finger nucleases^2,3^, and TALENs^4,5^, especially in the silk moth *Bombyx mori*, these approaches have resisted widespread application due to their laborious nature. We see two other main reasons manipulative genetics has failed to become routine in Lepidoptera. The first is that many lepidopterans are sensitive to inbreeding, and in some species it can be difficult to maintain experimental lines without special effort. The second is that lepidopterans appear to have an unusual resistance to RNAi^6,7^, a method that has dramatically accelerated work in many other groups of insects. Given this history of challenges in Lepidoptera, it is with great excitement that over the last few years we have seen an increasing number of studies that demonstrate the very high efficiency of CRISPR/Cas9-mediated genome editing in this group. Our own lab began experimenting with genome editing in butterflies in 2014, and we and our collaborators have now successfully edited over 15 loci across six species, generating both targeted deletions and insertions. The purpose of this review is to briefly summarize the current state of this fast-moving field and to provide practical advice for those who would like to use this technology in their own work.

## Published examples of Cas9-mediated genome editing in Lepidoptera

Between 2013 and early 2017 we identify 22 published studies applying CRISPR/Cas9 methods in Lepidoptera (Table 1). The earliest published reports of Cas9-mediated genome editing in Lepidoptera, from 2013 and 2014, all describe work done in *B. mori*^8-10^ – an experimental system that benefits from a large research community that had already developed efficient methods for injection, rearing, and genotyping. To our knowledge Wang et al. (2013)^8^ represents the first published report of Cas9-mediated genome editing in Lepidoptera, and set three important precedents. First, they established the protocol that has been more or less emulated by most following studies, where single guide RNAs (sgRNAs) are co-injected with Cas9 mRNA into early stage embryos. Second, they demonstrated that it is possible to co-inject dual sgRNAs to produce long deletions. In this respect, the 3.5 kb deletion they produced was an important early benchmark for demonstrating the possibility of generating long deletions in Lepidoptera. Third, they showed that deletions could occur in the germ line at a high enough frequency to generate stable lines.

**Table 1.**
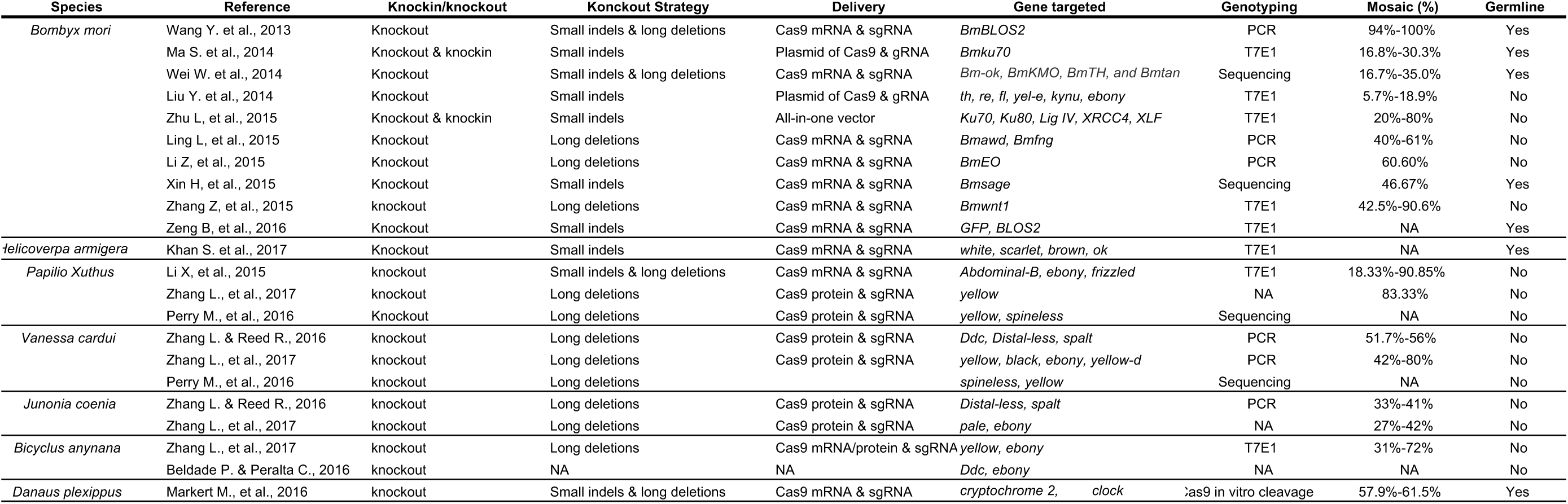
Comparison of CRISPR genome editing studies in Lepidoptera

After Wang et al. (2013)^8^ one of the next most important technical advancements came from Ma et al (2014)^9^, who showed that knockins could be achieved using a donor plasmid to insert a DsRed expression cassette using ~1kb homology arms. Following this, Zhu et al. showed successful epitope tagging of *BmTUDOR-SN* gene by CRISPR/Cas9 mediated knockin in *Bombyx* cells^11^. Unfortunately, to our knowledge, these remain the only two examples of lepidopteran knockins outside of the new data we present below. The first example of Cas9 genome editing in a species besides *B. mori* was described by Li et al. (2015)^12^, who produced deletions in three genes in the swallowtail butterfly *Papilio xuthus*. This was an important case study because it showed the general approach used by Wang et al. (2013)^8^ in *B. mori* could be transferred to other species and still retain the same level of high efficiency. Two more notable technical advancements include the production of an 18kb deletion in *B. mori* by Zhang et al. (2015)^13^ – the longest deletion we know of in Lepidoptera, and much longer than anything in *Drosophila* reports we have seen – and the direct injection of recombinant Cas9 protein instead of Cas9 mRNA^14,15^, which was an important improvement to the protocol that significantly simplifies the genome editing workflow.

Through our lab’s research on wing pattern development we have tried most of the methods described in the studies cited above, and we have gained significant experience in porting these protocols across species. We now perform targeted long deletions routinely and with a fairly high throughput. As of the end of 2016, we and our colleagues have successfully applied this general approach in six butterfly and two moth species (*Vanessa cardui, Junonia coenia, Bicyclus anynana, Papilio xuthus, Heliconius erato, Agraulis vanillae*, *B. mori*, and *Plodia interpunctella*), with each species requiring only minor modifications to physical aspects of egg injection protocol. As we describe below, we have also successfully achieved protein knockin tagging similar to Zhu et al.^11^, although our efficiency levels remain similarly low. In this report we outline the approach that we have found to be the most time-and cost-efficient and transferable between species (Fig. 1A).

**Figure 1.**
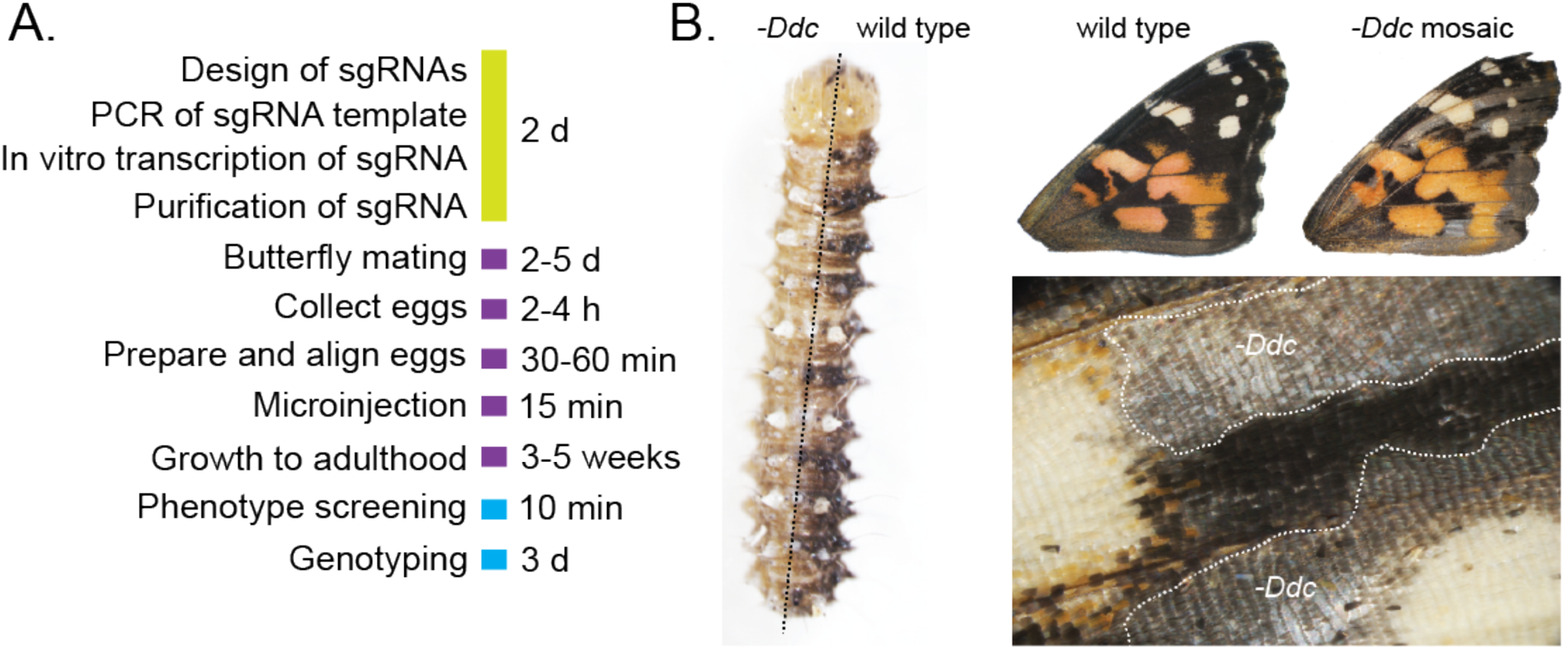
Timeline and example outcome of G_0_ CRISPR/Cas9 mosaic knockout experiments in butterflies. (A) Overview and timeline of mutant generation by CRISPR/Cas9 injection to butterfly embryos. (B) Example of larval and adult wing somatic mosaic phenotypes resulting from knockout of the melanin pigmentation gene *Ddc.*

## Experimental Design

### Deletions

Loss-of-function deletion mutations can be generated by non-homologous end joining (NHEJ) following double strand breaks (DSBs). Both small indel (single cleavage) and long deletion knockout strategies (co-injection of two sgRNAs) have been employed in Lepidoptera (Table 1). Our lab currently favors long deletions using dual sgRNAs because it facilitates rapid screening and genotyping of mutants using PCR and regular agarose gel electrophoresis. Small indels produced by single cleavages are too small to detect easily using normal agarose gels. Dual sgRNA deletions, however, can be tens, hundreds, or thousands of base pairs long and are easy to identify in gels. sgRNA target sites can be easily identified simply by scanning the target region for GGN_18_NGG or N_20_NGG motifs on either strand using the CasBLASTR web tool (http://www.casblastr.org/). In our experience the relative strandedness of sgRNAs does not appear to have a significant effect on the efficiency of double sgRNA long deletion experiments. If a reference genome is available candidate sgRNA sequences should be used for a blast search to confirm there are not multiple binding sites that may produce off target effects. The injection mix we typically use is 200 ng/μl Cas9 and 100 ng/μl of each sgRNA – this will tend to give larger effects and is suitable for less potentially lethal loci. For targets that may result in more deleterious effects we recommend decreasing the amount of Cas9/sgRNA mix and injecting later in embryonic development to induce fewer and smaller clones. We have been able to induce mosaic mutants (e.g. Fig. 1B) using as low as 20 ng/μl Cas9 and 50 ng/μl of each sgRNA in different butterfly species.

### Insertions

CRISPR/Cas9-induced site specific DSBs can be precisely repaired by homology directed recombination repair (HDR). The HDR pathway can replace an endogenous genome segment with a homologous donor sequence, and can thus be used for knockin of foreign DNA into a selected genomic locus. To our knowledge there are only two published examples of this approach in Lepidoptera, both of which are in *B. mori*^9,11^. To test the feasibility of this approach in butterflies we sought to insert an in-frame EGFP coding sequence into to the *V. cardui dopa decarboxylase* (*Ddc*) locus using a donor plasmid containing the EGFP coding sequences and homologous arms matching endogenous sequences flanking the Cas9 cut sites (Fig. 2A). As shown in Fig. 2B, EGFP fluorescence was detected in clones in the mutant caterpillars. As well, PCR analysis with primers flanking the 5′ and 3′ junctions of the integration showed a clear band in mutants, but not in wild type (Fig. 2C). Our results show that donor DNA with ~500 bp homology arms is sufficient for precise in-frame knockins. Compared to NHEJ-mediated high efficiency knockouts (69% in the case of *V. cardui Ddc* deletions knockouts^14^), the rate of HDR mediated targeted integration is low, at ~3% in our most recent trials. It has been shown that knocking out of factors in the NHEJ pathway can enhance the HDR pathway and increase gene targeting efficiency in *Bombyx*^9,11^. Some Cas9 mediated homology-independent knockin approaches have shown higher efficiency rates in zebrafish^16^ and human cell lines^17^, suggesting NHEJ repair may provide an alternate strategy to improve incorporation of donor DNA in Lepidoptera.

**Figure 2.**
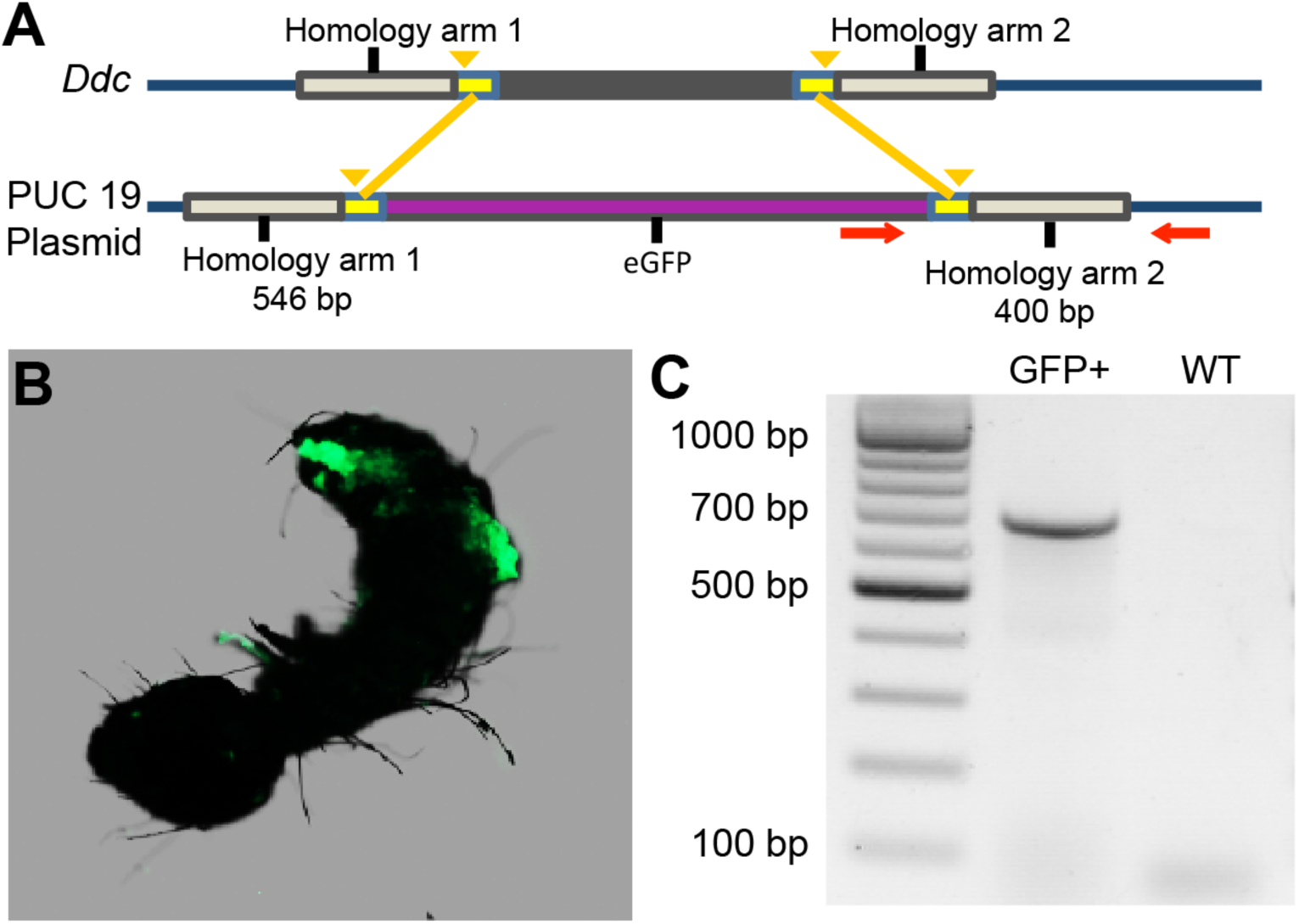
Knockin tagging of the *Ddc* gene in *V. cardui*. (A) Schematic overview of the *Ddc* locus and donor construct consisting of homology arms, EGFP coding region, and genotyping primers. PAM regions are marked by yellow, cut sites are marked by yellow arrowhead, and genotyping primers are marked by red arrows. (B) Strong mosaic EGFP expression in a knockin caterpillars visualized by fluorescent microscopy. (C) PCR analysis demonstrates using the primers in (A) show the insertion of EGFP into the *Ddc* coding region.

## Embryo Injection

When we adapt CRISPR/Cas-9 genome editing to a new species the greatest technical challenge we face typically lies in optimizing the injection protocol. The primary reason for this is that the eggs of different species can be quite different in terms of how difficult they are to puncture with a glass needle and how they react to mechanical injection, especially in terms of internal pressure and post-injection backflow.

### Injection Needles

Proper needle shape is critical for achieving successful egg injections in Lepidoptera. In our experience some taxa like *Heliconius spp.* have very soft, easy-to-inject eggs that present very few problems, and are more robust to variation in needle shape. Many lepidopterans, however, have difficult-to-puncture eggs that have high internal pressure. The key challenge for these eggs is to use needles that are strong enough to penetrate tough eggshells, but are not so wide as to weaken pressure balance or destroy embryos. For instance, needles that are too long and narrow can break easily when used on tough eggs, and will clog at a high frequency. Conversely, needles that have a very wide diameter will tend to have problems with pressure loss and backflow. We recommend the needle shape shown in Fig. 3A which is characterized by a short rapid taper to a fine point. We have found that this shape provides enough strength to puncture fairly tough eggshells, yet is relatively resistant to clogging and pressurization problems. Our initial attempts at pulling needles like this with a traditional gravity needle puller failed. We currently pull our needles using a velocity sensitive Sutter P-97 programmable needle puller, which works very well for crafting nuanced needle shapes. We currently prefer to use Sutter Instrument 0.5mm fire polished glass capillary needles (Sutter BF-100-50-10) and 3 mm square box heating filaments (Sutter FB330B). Although settings will vary by instrument and filament, we use a single cycle program on our puller with parameters HEAT 537, PULL strength 77, VELOCITY (trip point) 16, and TIME mode (cooling) 60. Among these parameters, the HEAT value has to be adjusted relative to the RAMP value, which is specific to certain instruments – different pullers can produce slightly different needle shapes even with the same parameter settings. We provide our settings as a starting point for other users to work towards optimizing production of needles with a steep taper and a large orifice as shown in Fig. 3A.

**Figure 3.**
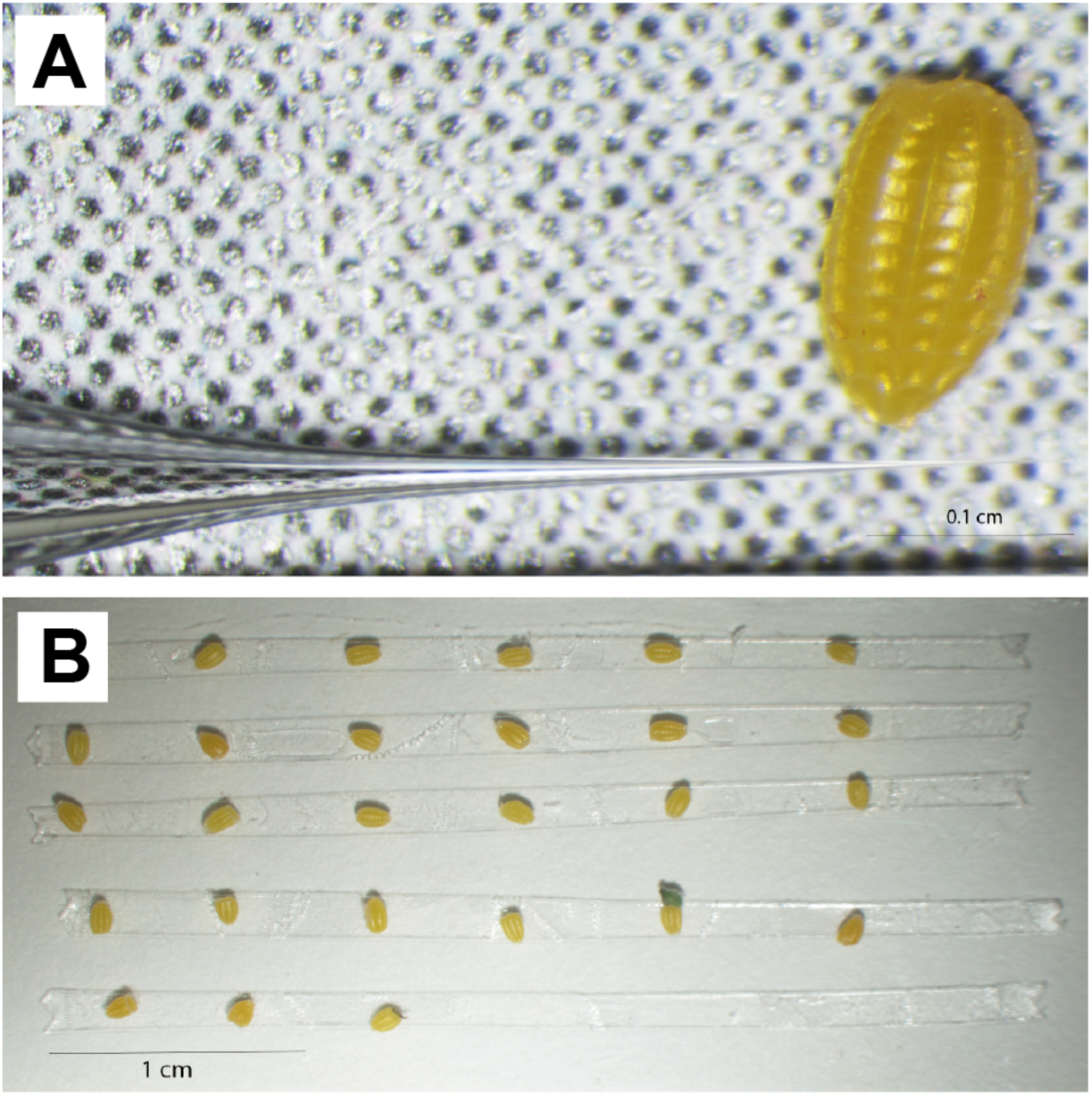
Needle shape and egg arrangement for butterfly embryo injections. (A) The injection needle shape we prefer has a steep taper and a relatively large orifice. Here a preferred needle is shown next to a *Heliconius* egg. (B) An example of arranging *Heliconius* eggs on double-sided tape on a microscope slide just before injection.

### Egg Treatment

Egg treatment is different among eggs from different taxa. For butterflies with soft eggs such as *Heliconius*, *Agraulis*, and *Danaus*, freshly collected eggs can be immediately arranged on double-sided adhesive tape on a microscope slide (Fig. 3B) and injected. For those eggs with a relatively soft chorion but high pressure, like *V. cardui*, collected eggs should be arranged on a slide, and then kept in a desiccation chamber for 15min before injection. We use a simple sealed petri dish filled with desiccant for this purpose. For species with thick-shelled eggs like *J. coenia*, we recommend that eggs be dipped in 5% benzalkonium chloride (Sigma-Aldrich, St Louis, MO, USA) for 90s to soften the chorion, and then washed in water for 2 min before mounting on microscope slide. We also tried treatment with 50% bleach solution to soften eggs, however this significantly reduced the hatch rate. Softened eggs can then be dried in a desiccation chamber for 15 min and injected.

### Injection Timing

In all published cases we are aware of injections of sgRNA and Cas9 (either mRNA or recombinant protein) were completed between 20 min and 4 hrs after oviposition, when embryos are presumed to be in an acellular syncytial state. Most of our injecting experience has been in eggs 1-3 hrs old. Although we have not rigorously quantified this effect, after extensive work with pigmentation genes in *V. cardui* we found that injecting earlier (e.g. at 1 hr) typically produces more and larger mutant clones compared to injection performed later (e.g. at 4 hrs). This is consistent with previous studies that have found a higher deletion frequency when embryos are injected at earlier versus later stages^12^. Thus, for most of our deletion experiments we aim to inject ~1-2 hrs after oviposition. If we expect that deletion of the locus will have a strongly deleterious or embryonic lethal effect, we will begin by injecting at 2-4 hrs to decrease the magnitude of somatic deletions.

### Egg Injection

The key concern during injection is to minimize damage as much as possible. An optimum angle for needle insertion is about 30-40 degrees in our experience. We prefer to use a Narishige MM-3 micromanipulator for full three-dimensional control of the needle during injection. In the butterfly species we have worked with the location of the injection does not seem to have a major impact on editing efficiency, although in *V. cardui* we get a slightly higher survival rate by injecting into the side near the base of the egg. Proper positive balance pressure is critical for successful injection. Users should adjust balance pressure to a point where the needle is just able to retain the solution. Prior to any egg injection, adjust the injection pressure and time to ensure the flowing droplet is visible when pressing the injector’s footswitch. We have worked extensively with two different injectors: a Harvard Apparatus PLI-100 Pico-Injector and a Narishige IM 300 Microinjector. In our experience PLI-100 Pico-Injector has better sensitivity in terms of balance pressure, which is very important for species with high pressure eggs like *V. cardui* and *J. coenia*. The IM 300 does not perform as well on these eggs. The other two injectors we know of that also work well for butterfly eggs are Eppendorf FemtoJet microinjector and Drummond Nanoject III. In our hands, we use 10 psi injection pressure and 0.5 psi balance pressure for soft-shelled eggs with the Narishige IM 300 injector, and 20 psi injection pressure and 0.8 psi balance pressure for *V. cardui* and *J. coenia* eggs with the PLI-100 Pico-Injector. After injection we maintain slides with the injected eggs in a petri dish and move larvae to their rearing containers immediately upon emergence.

## Interpreting somatic mosaics

While several studies have been published that describe the germ line transmission of edited alleles in *B. mori*, thus far most studies in Lepidoptera have focused on interpreting deletion phenotypes in G0 somatic mosaics. Maintaining edited genetic lines is necessary for looking at the homozygous effects of specific edited alleles, and will also be essential for a future generation of more sophisticated knockin studies. Maintaining edited lines presents a few challenges in Lepidoptera, however. First, the deletion phenotypes of many interesting genes would likely be embryonic lethal. For example, our lab has thus far been unsuccessful in efforts to produce living larvae with *wingless* or *Notch* coding region deletions, which is unsurprising because these genes are known to be essential for early embryonic development in insects. For loci like these we can confirm deletions by PCR and sequencing, but all embryos with deletions die before or shortly after hatching. Second, based on our experience with inbreeding attempts in *Heliconius spp.*, *V. cardui*, and *J. coenia*, and through discussions with colleagues working in other systems, it is clear that many Lepidoptera are sensitive to inbreeding and lines will die out quickly unless fairly large stocks are kept. Large stocks then make it more difficult to identify individuals with specific genotypes. While maintaining lines is certainly possible in many lab-adapted species, it can require a fair amount of time and resources.

Because of the challenges posed by the embryonic lethality of many target genes, along with the difficulty of maintaining and genotyping edited lines, most of our attention has focused on analysis of mosaic G_0_ phenotypes. One obvious advantage of focusing on somatic mosaics is that data can be collected in a single generation. Another advantage is that the phenotypic effects of lesions are limited to the subset of cell lineages (clones) hosting deletion alleles, thus reducing the deleterious effects of many deletions. Because of their clear phenotypes, knockout work on melanin pigmentation genes has allowed a very useful visual demonstration of the nature of somatic mosaicism in injected animals. Our work on eight pigmentation genes across several butterfly species^18^ has allowed us some general insights into work with mosaics. First, as described above, we found a loose association between the number and size of clones and the timing of injection, where earlier injections with higher concentrations of Cas9/sgRNA complexes tend to produce larger clones. We have not attempted to quantify this effect, but across replicated experiments our tentative conclusion is that this is a real and consistent phenomenon. This is important because it gives rough control over the strength of a phenotype, and can thus be important for trying to get small non-deleterious clones for an otherwise lethal gene. Conversely, by injecting at very early stages to knock out minimally pleiotropic genes we could produce animals with very large clones, such as entire wings.

One challenge of working with somatic mosaic lies in detecting and interpreting more subtle phenotypes. Most of the phenotypes published to date, such as loss of wing pattern features like eyespots or production of discolored patches, are fairly obvious and/or far outside the range of natural variation. Without having a dramatic phenotype or independent clone boundary marker, however, minor or highly localized effects can be difficult to differentiate from natural variation. It is possible that quantitative image analysis approaches could address this issue, although we are unaware of published examples of this. In our own work we have relied on two main criteria to validate putative deletion phenotypes: (1) Replicates, which, of course, are useful for increasing confidence (we typically aim for a minimum of three, although the number of replicates required to make a particular inference is somewhat arbitrary and there is no standard), and (2) asymmetry, which is perhaps the most powerful criterion for inferring more subtle deletion phenotypes. Because natural variation is ordinarily symmetrical, strongly asymmetric phenotypes are best explained by left/right variation in clonal mosaics.

## Genotyping

To validate that genome editing is occurring as expected at the appropriate locus, it is necessary to perform genotyping on experimental animals. We have found that the simplest and most robust genotyping approach is to design PCR primers flanking deletion sites, and then to compare PCR product sizes between wild type and experimental animals. We recommend that genotyping amplicons cover less than 1.5 kb and be at least 100 bp outside of the closest sgRNA site to allow proper band size resolution and detection of large deletions. This approach works best for long deletions produced by double sgRNAs – indels induced by repair of a single Cas9 cut site will usually be too small to detect by PCR alone. For this reason, our lab always uses double sgRNAs to produce deletions. These PCR products can also be cloned and sequenced for further validation, as well as to better characterize the diversity and nature of deletion alleles. If a single sgRNA is used it is likely that deletion alleles will need to be sequenced to confirm lesions. To genotype insertions PCR primers flanking the insertion site may be used similarly, or one may also use a primer inside the transgene (e.g. Fig 2C).

A current challenge in genotyping edited animals is the lack of tools to rigorously confirm specific deletion alleles in specific cell populations. First, there is the physical problem of isolating a population of cells representing a single pure clone. To our knowledge, this has not been done in insects outside of using transgenic cell sorting methods^19^. Even carefully dissected presumptive clones cannot be assumed to be pure clonal cell populations. Indeed, to our knowledge there is not yet a practical method developed to firmly associate specific alleles with specific phenotypes. This challenge also makes it difficult to decisively confirm whether a clone is monoallelic (i.e. has a single edited allele) vs. biallelic (i.e. has two edited alleles), thus making it difficult to infer dosage effects without additional information. Therefore, even though some previous studies present DNA sequences of edited alleles isolated from tissues including cells with deletion phenotypes (e.g. whole embryos), none of these studies rigorously associate individual alleles with specific clones because it cannot be ruled out that the tissue samples maybe have contained multiple monoallelic or bialleleic clones. A second challenge for genotyping specific clones is that some tissues of special interest, such as adult cuticle structures, including wing scales, do not have genomic DNA of sufficient quality to permit straightforward PCR genotyping, especially for longer amplicons. Thus, even if methods become available for isolating specific clone populations, there will still be limitations when dealing with some tissue types. Given the challenges outlined above, readers should understand that most genotyping to date should be seen as a validation of the experimental approach (editing accuracy and efficiency), and not necessarily as completely decisive confirmation that specific alleles underlie a certain clone phenotype.

## Future prospects

CRISPR/Cas9 genome editing is rapidly revolutionizing genetic work in Lepidoptera, as it is across all of biology. It is now fairly straightforward to quickly and cheaply induce long, targeted deletions in virtually any species that can be reared in captivity. Published reports to date have focused on producing deletions in gene coding regions, however we anticipate there will be significant interest in also applying long deletion approaches to test the function of non-coding regulatory regions, especially now that cis-regulatory elements can be functionally annotated with high resolution thanks to methods like ChIP-seq^20^. Pilot work shown here and elsewhere also shows that targeted insertions are possible as well, thus promising even further developments on the near horizon such as protein tagging, reporter constructs, and tissue-specific expression constructs. Right now the main challenge with knockin strategies is the relatively low efficiency rate, although newer technologies such as non-homologous end joining mediated knockin^16^ promise to dramatically improve this. Perhaps the most exciting thing about CRISPR-associated genome editing approaches, though, is the straightforward portability of the technology between species. This is truly an exciting time to be a comparative biologist.

## Acknowledgments

We thank Arnaud Martin for extensive discussions regarding genome editing methods in butterflies, Joseph Fetcho for early assistance with needle pulling and injection procedures, and Katie Rondem for helpful comments on the manuscript. This work was funded by United States National Science Foundation grant DEB-1354318

## Appendix A detailed example of CRISPR/Cas9 genome editing in the painted lady butterfly *V. cardui*

The following procedure provides guidelines to generate genomic deletions in the butterfly *V. cardui* using the CRISPR/Cas9 nuclease system. This protocol includes a specific example of the Reed Lab’s work deleting the melanin pigmentation pathway gene *Ddc* as previously reported^14^.

### 1. Target design

No genome reference was available for *V. cardui* when we first began our experiment, so we used a transcriptome assembly^18^ to identify sequences of the *Ddc* coding region. Primers GCCAGATGATAAGAGGAGGTTAAG and GCAGTAGCCTTTACTTCCTCCCAG were designed to amplify and sequence the target region of the genome, and exon-intron boundaries were inferred by comparing genomic and cDNA sequences. We recommend designing target sites at exons because they are more conserved than introns and therefore provide more predictably consistent matches between sgRNAa and genomic targets. We design sgRNAs by scanning for GGN_18_NGG or N_20_NGG pattern on the sense or antisense strand of the DNA. Target sequences GGAGTACCGTTACCTGATGA**<u>AGG</u>** and **<u>CCT</u>**CTCTACTTGAAACACFACCA (PAM sequences underlined) were designed to excise a region of 131 bp spanning the functional domains of the DDC enzyme. sgRNA oligos containing T7 promoter, target sequences and sgRNA backbone were synthesized by a commercial supplier (Integrated DNA Technologies, Inc.). Of note, the PAM sequence is not included in the CRISPR forward primer.

CRISPR forward oligos:

*Ddc sgRNA 1*: GAAATTAATACGACTCACTATA**GGGATCAGCTTTCGTCTGCC**GTTTTAGAGCTAGAAAT AGC

*Ddc sgRNA2*: GAAATTAATACGACTCACTATA**GGAGTACCGTTACCTGATGA**GTTTTA GAGCTAGAAATAGC

CRISPR universal oligo: AAAAGCACCGACTCGGTGCCACTTTTTCAAGTTGATAACGGACTAGCCTTATTTTAACTTG CTATTTCTAGCTCTAAAAC

### 2. sgRNA production

#### 2.1 sgRNA template generation

- With the oligos generated in the preceding step, use High-Fidelity DNA Polymerase PCR Mix (NEB, Cat No. M0530) to generate the template for each sgRNA with CRISPR forward and reverse oligos. We recommend using DEPC-free nuclease-free water (Ambion, Cat No. AM9938).

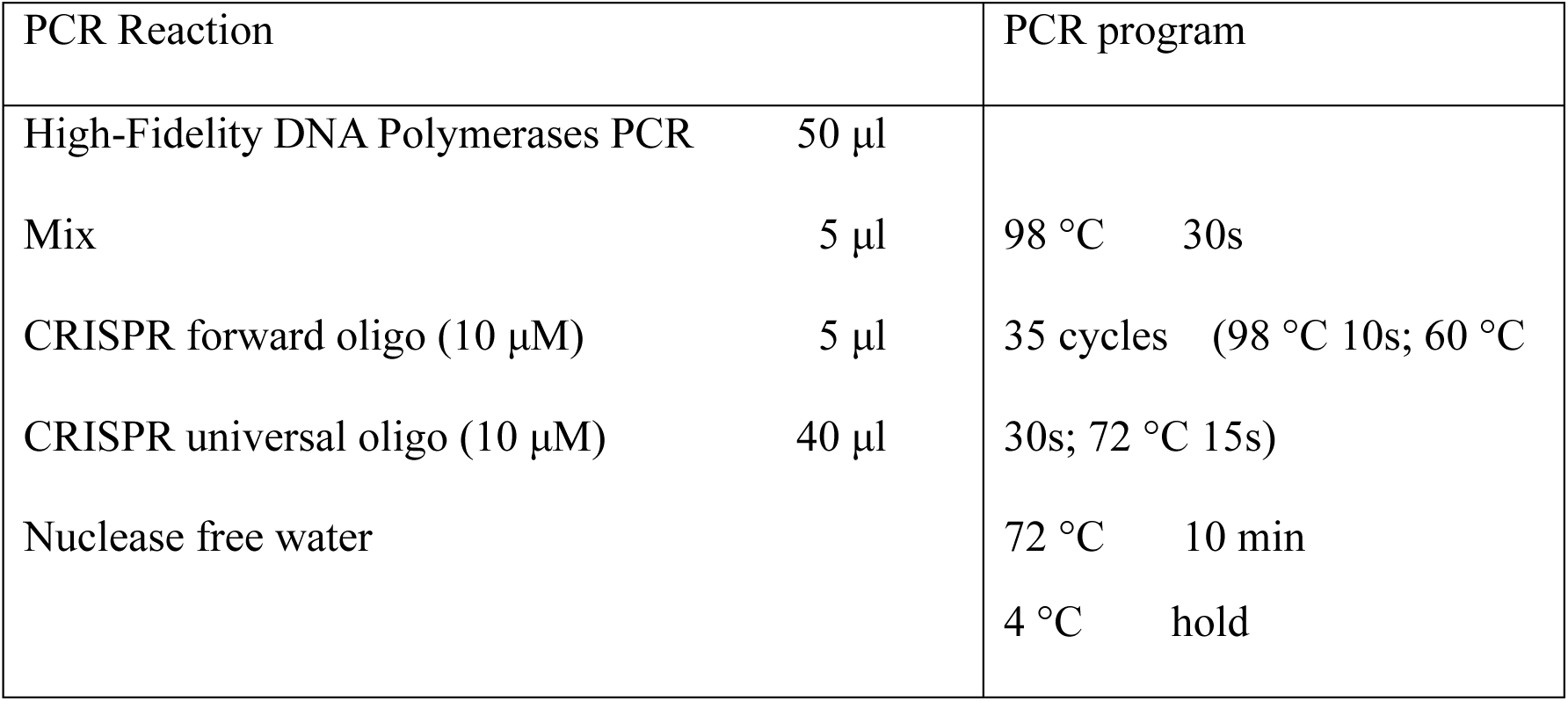

- Purify the PCR reaction with MinElute PCR purification kit (Qiagen, Cat No. 28004) following the kit instructions and eluting in 15 μl nuclease free water.
- Dilute 1μl of this reaction with 9 μl nuclease free water, then run on a gel and a fluorometer (e.g. Qubit) to confirm purity, integrity, fragment length, and yield. It is also possible to use gel extraction at this stage if non-specific products are present.
- The expected size should be around 100bp, and the expected yield should be around 200 ng/ul.

#### 2.2 in vitro transcription (IVT)

- Generate sgRNAs by in vitro transcription of the sgRNA PCR template using the T7 MEGAscript kit (Ambion, Cat. No. AM1334). When producing and handling RNA, it is important to wear gloves, and clean equipment and benches with detergent prior to use to avoid RNAase contamination. Pipette tips with filters can also be beneficial to prevent contamination from pipettes.

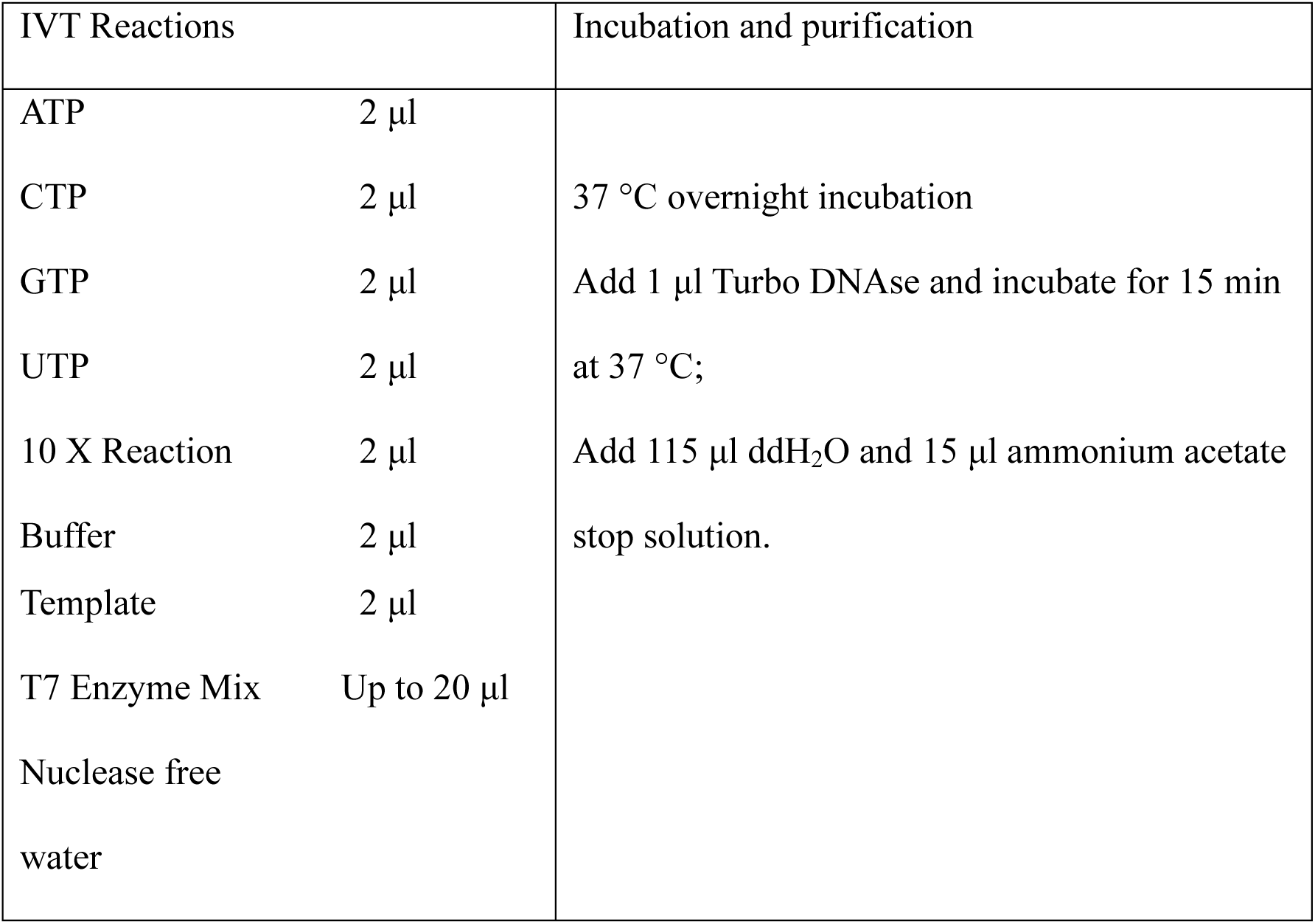

- Extract sgRNA by adding 150 μl phenol:chloroform:isoamyl alcohol (25:24:1) at pH 6.7 (Sigma-Aldrich, Cat No. P2069), and vortex thoroughly for 30 s.
- Separate phases by centrifugation at 10,000 x g for 3 min at room temperature, and remove the upper phase to a fresh tube.
- Precipitate the RNA by addition of an equal volume (150 μl) of cold isopropanol (Sigma-Aldrich, Cat No. I9516).
- Mix thoroughly, and incubate at -20 °C for greater than 2 hours (can be left overnight).
- Collect RNA by centrifugation at 17,000 x g for 30 min at 4 °C.
- Wash pellet twice in 0.5 ml room temperature fresh made 70% ethanol, centrifuging at 17,000 x g for 3 min at 4 °C between each wash.
- Remove the remaining liquid and dry RNA pellet for 3 min at room temperature.
- Resuspend in 30 ul nuclease-free water.
- Measure concentration on a Qubit. The expected concentration should be around 2 μg/ul; sgRNAs can be stored at -80 °C.
- MEGAclear^™^ Transcription Clean-Up Kit (Ambion, Cat No. AM1908) also works very well for sgRNA purification.

### 3. Cas9 production

Cas9 is typically provided by injection of a plasmid, mRNA, or recombinant protein. We have tried both Cas9 mRNA and protein injections, and both yield similarly efficient mutation rates in butterflies. However, we recommend using commercially available Cas9 protein (PNA Bio, Cat No. #CP01) because it is more stable than Cas9 mRNA and is easier and faster to use.

- Cas9 mRNA is generated by in vitro transcription of the linearized MLM3613 (Addgene plasmid 42251) plasmid template. The mMessage mMachine T7 kit (Ambion, Cat No. AM1344) is used to perform in vitro transcription with T7 RNA polymerase, followed by in vitro polyadenylation with the polyA tailing kit (Ambion, Cat No. AM1350). An Agilent Bioanalyzer, or similar instrument, should be used to check the size and integrity of Cas9 mRNA. Note that Cas9 mRNA can show some degree of degradation yet still produce fairly efficient results.

### 4. Egg injection and survivor ratio calculation

- Collect eggs for 2-4 hours by placing a host plant leaf into the butterfly cage.
- For thick chorion eggs (e.g. *J. coenia*), dip eggs in 5% benzalkonium chloride for 90 s.
- Cut double-sided tape into several thin strips and fix them to a glass slide.
- Using a paintbrush line the eggs onto the double-sided tape.
- For high-pressure eggs (e.g. *V. cardui* or *J. coenia*) place the slide in a desiccation chamber for 15 min before injection.
- Mix Cas9 and CRISPR sgRNAs prior to microinjection.

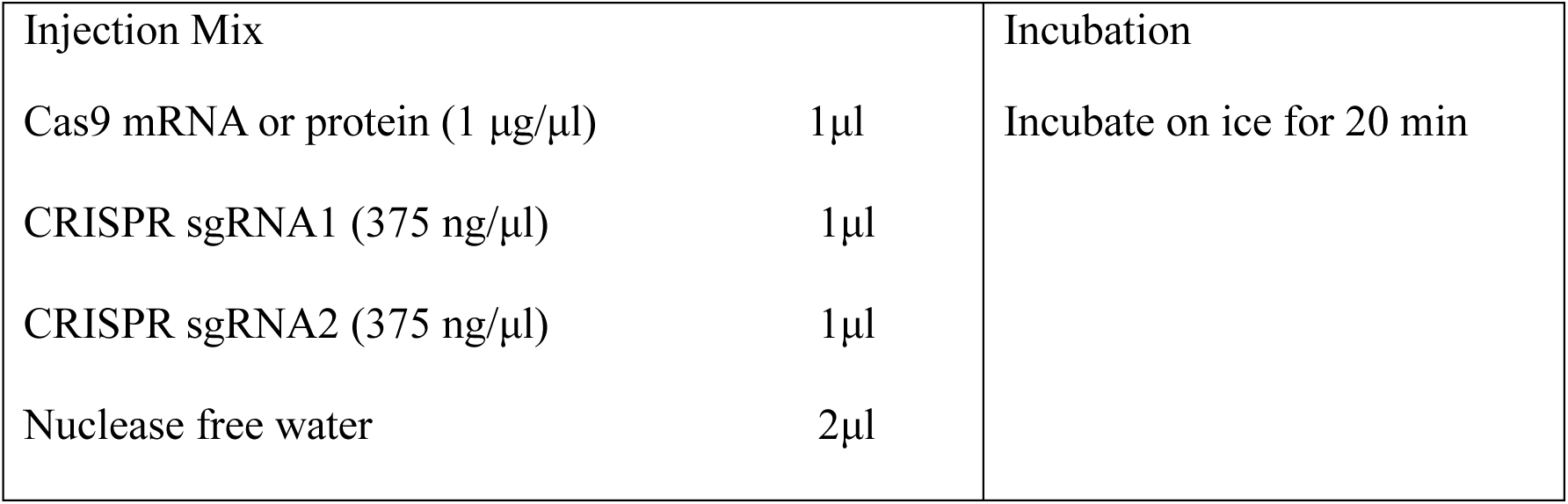

- Break the closed tip of the needle with an optimum angle about 30-40 degree.
- Load the needle with 0.5 μl injection mix by capillary action, or by using by Eppendorf^™^ Femtotips microloader tips (Eppendorf, Cat No. E5242956003).
- One-by-one inject the eggs with the injector.
- Generally, higher amounts of sgRNA and Cas9 protein will increase mutation rate and decrease egg survival (hatch rate).

### 5. Genotyping for modification

- In order to investigate the efficiency of CRISPR/Cas9 mediated *Ddc* knockout, we randomly surveyed 81 first instar caterpillars. DNA was extracted according to Bassett et al.^28^ to confirm CRISPR/Cas9 lesions. Generally, place one caterpillar in a PCR tube and mash the caterpillar for 30 s with a pipette tip in 50 μl of squishing butter (10 mM Tris-HCl, pH 8.2, 1mM EDTA, 25 mM NaCl, 200 μg/ml proteinase K). Incubate at 37 °C for 30 min, inactivate the proteinase K by heating to 95 °C for 2 min and store in -20 °C for PCR genotyping. Genotyping can also be done with adult butterfly leg DNA by using proteinase K in digestion buffer. We typically use QIAamp DNA mini Kit (Qiagen, Cat No. 51304) for DNA extraction when genotyping from muscle tissue.
- Design genotyping primers outside of the target region. For *Ddc*, genotyping forward (GCTGGATCAGCTATCGTCT) and reverse primers (GCAGTAGCCTTTACTTCCTCCCAG) were designed and gave a 584bp PCR fragment in the wild type individuals.
- Mix PCR reagents. PCR fragments containing two sgRNA target sites are expected to produce smaller mutant bands than wild type.

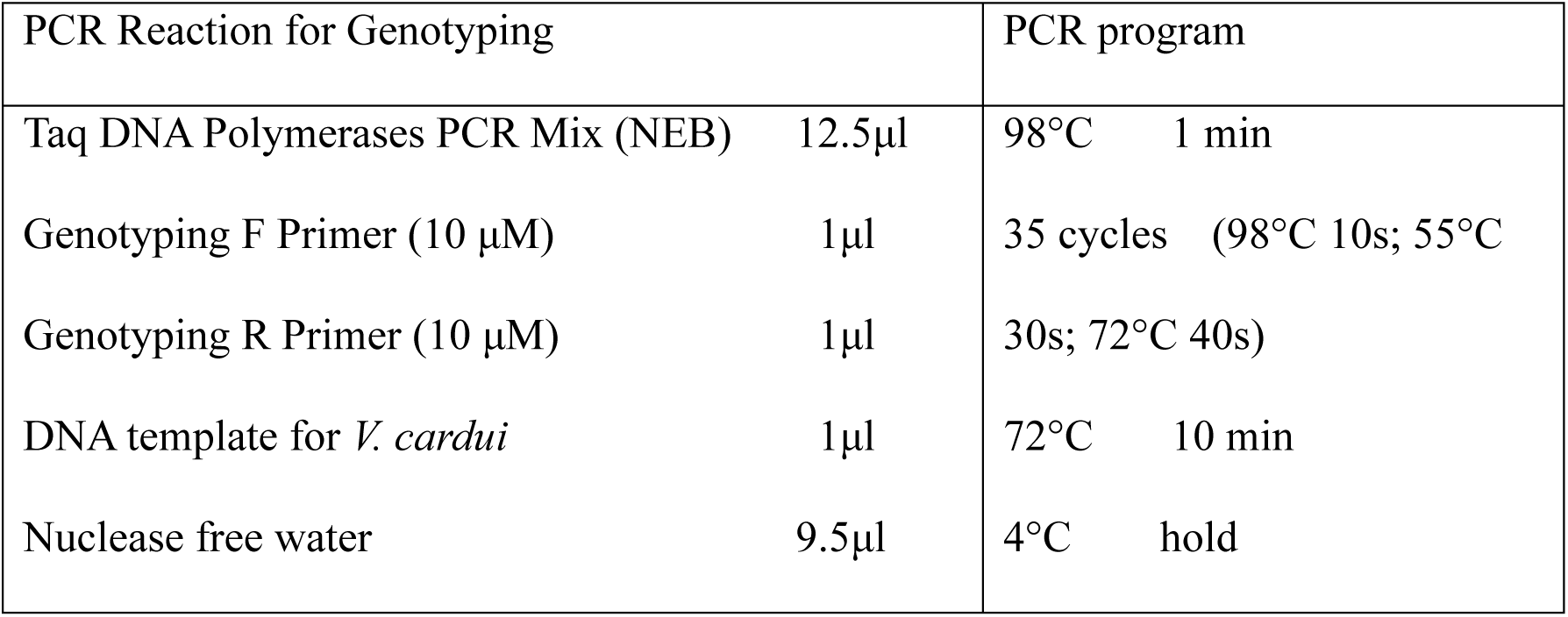

- Recover mutant bands by gel-extraction using Minelute gel extraction kit (Qiagen, Cat No. 28604).
- Ligate recovered DNA fragment to T4 vector for TA-cloning using a TA cloning kit (Invitrogen, Cat No. K202020).
- Extract plasmid with mutant DNA fragment using QIAprep Miniprep Kit (Qiagen, Cat No. 27104).
- Sequence plasmids and align mutant sequences to wild type sequences to confirm deletions. (Figure 1a in Zhang & Reed, 2016)^14^.

